# Exact Distribution of Linkage Disequilibrium in the Presence of Mutation, Selection or Minor Allele Frequency Filtering

**DOI:** 10.1101/794347

**Authors:** Jiayi Qu, Stephen D Kachman, Dorian Garrick, Rohan L Fernando, Hao Cheng

## Abstract

Linkage disequilibrium (LD), often expressed in terms of the squared correlation (*r*^2^) between allelic values at two loci, is an important concept in many branches of genetics and genomics. Genetic drift and recombination have opposite effects on LD, and thus *r*^2^ will keep changing until the effects of these two forces are counterbalanced. Several approximations have been used to determine the expected value of *r*^2^ at equilibrium in the presence or absence of mutation. In this paper, we propose a probability-based approach to compute the exact distribution of allele frequencies at two loci in a finite population at any generation *t* conditional on the distribution at generation *t* − 1. As *r*^2^ is a function of this distribution of allele frequencies, this approach can be used to examine the distribution of *r*^2^ over generations as it approaches equilibrium. The exact distribution of LD from our method is used to describe, quantify and compare LD at different equilibria, including equilibrium in the absence or presence of mutation, selection, and filtering by minor allele frequency. We also propose a deterministic formula for expected LD in the presence of mutation at equilibrium based on the exact distribution of LD.

## Introduction

Linkage disequilibrium (LD), the nonrandom association of alleles at two or more loci, is an important concept in various areas of genetics, including evolutionary, quantitative and statistical genetics. In evolutionary genetics, LD is used to detect the genomic locations of historical selection and to estimate the time of divergence and geographic subdivision between populations (Sved and Hill 2018). For example, LD was used to date the divergence of the European population from the African population using the HapMap data (Sved *et al.* 2008). In the field of quantitative and statistical genetics, methods for genomic prediction and genome-wide association studies (GWAS) rely on the existence of LD between the molecular markers and the unobserved quantitative trait loci (QTL).

Two statistics that have been widely used to quantify the LD between allelic values at two loci are: their covariance (*D*); or their squared correlation (*r*^2^) (Hill and Robertson 1968). For a diallelic system, they can be expressed as

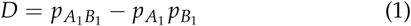

and

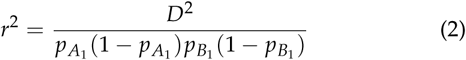

respectively, where 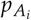 represents the frequency of the *ith* allele at locus 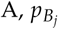 represents the frequency of the *jth* allele at locus B, and 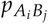 represents the frequency of haplotype *A*_*i*_*B*_*j*_ in the population. Although the covariance of different pairs of alleles depends on their corresponding haplotype frequencies and the respective allele frequencies, a single value of D is sufficient to characterize the disequilibrium between two loci in a diallelic system, i.e., 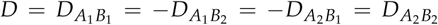. The value of *D* depends on how the alleles are coded (i.e., alleles *A*_1_ and *A*_2_ could be coded as 0 and 1 or as 1 and 0) while the value of *r*^2^ is invariant to this choice. Therefore, *r*^2^ is increasingly used as a metric to quantify LD in the literature.

In a finite population, allele frequencies at two loci are subject to random fluctuations due to the stochastic sampling of a finite number of gametes across generations. This random change of the gene frequency between one generation and the next is genetic drift. Due to the randomness of the genetic drift, one initial random-mating population can evolve into one of a finite collection of sub-populations or lines with different allele frequencies at the two loci, and thus with different LD. Usually, this finite collection of possible lines is jointly considered to understand the distribution of the allele frequencies and LD of two linked loci over generations. This is equivalent to considering a finite collection of pairs of loci in one line. The consequences of the genetic drift, mutation, and selection for the collection of lines of two loci equally apply to a collection of pairs of loci in one line. The former conceptualization is used in this paper to understand the effects of genetic drift, mutation, selection and minor allele frequency (MAF) cutoff on the distribution of LD.

Generally, in a finite population, the opposite effects on LD of genetic drift and recombination lead to an equilibrium value for the expectation of *r*^2^ when these two forces are counterbalanced. It has been shown that for large values of *N*_*e*_*c*, the equilibrium value for the expectation of 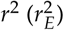 is approximately:

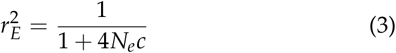

where *c* is the recombination rate and *N*_*e*_ is the effective population size (Sved 1971; Sved and Feldman 1973; Hill 1975).

The above formula, however, does not consider the variation introduced by mutation in the population. When mutation is considered, a balance is expected between the loss of variation by the fixation of one or other allele, and the replenishment of an extinct allele by mutation. An approximation for the equilibrium value of *r*^2^ in the presence of mutation in a finite population has been developed in Ohta and Kimura (1971) and Hill (1975). Instead of using *E*(*r*^2^), they used a quantity 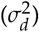 named the standard linkage deviation (Ohta and Kimura 1969) to describe the status of this equilibrium, where 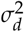 is an approximation of the expected value of *r*^2^ expressed as the ratio of the expectations of the numerator and the denominator of *r*^2^ (Ohta and Kimura 1971):

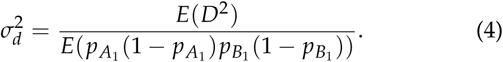

The equilibrium value of 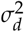 is approximated as:

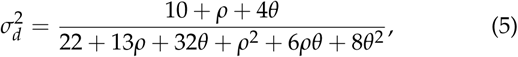

where *ρ = 4N*_*e*_*c, θ = 4N*_*e*_*u* and *u* is the mutation rate per site (Ohta and Kimura 1971). The equilibrium value of 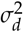 as an approximation of 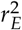 has been described in Walsh and Lynch (2018). Similarly, this value is considered in this paper as an approximation of 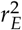 in the presence of mutation.

Selection is another important force affecting LD that was not considered in either Sved’s or Hill’s formulas. Selection occurs at causal variants or QTL during evolution (natural selection) or selective breeding (artificial selection). The impact of selection can either decrease or increase the LD in a population given different scenarios (Mitchell-Olds *et* al. 2007). The equilibrium among mutation and selection is of special interest in terms of the distribution of allele frequencies and of *r*^2^.

In contrast to the approximate methods mentioned above, in this paper we will derive the exact distribution of allele frequencies and LD of two linked loci at equilibrium in the absence or presence of mutation, selection and MAF cutoff. The distribution of LD in the presence of filtering by MAF is considered in our study due to the prevalent practice of filtering out marker loci with low MAF in genomic analyses used for genetic evaluation or QTL discovery (GWAS).

Our exact distributions are first used to validate the approximate deterministic formulas for 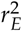, such as Sved’s and Hill’s approximations. Next we describe, quantify and compare 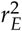 at different equilibria: including equilibrium in the presence of mutation; equilibrium in the presence of mutation and selection; with or without filtering by MAF. Finally, we calibrate Sved’s deterministic formula for 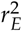 in the presence of mutation based on the exact distribution of LD. The objective of this paper is to present a computational approach to derive the exact distribution of LD over generations and use it to study the distribution of LD whether or not there is mutation, selection, or filtering by MAF.

## Materials and Methods

In this section, we will show how to compute the exact distribution of allele frequencies at two linked loci in generation *t*, given their distribution in generation *t* − 1, as a function of the effective population size, the recombination rate between the loci, the mutation rate, the selection coefficient, and the MAF threshold. As LD is a function of the allele frequencies at the two loci, its distribution over generations can be calculated based on the distribution of the allele frequencies.

### Transition-Matrix Approach in the Presence of Mutation, Selection or Filtering by Minor Allele Frequency

Two diallelic loci *A* and *B*, with alleles *A*_1_ and *A*_2_ at the *A* locus and alleles *B*_1_ and *B*_2_ at the *B* locus, are considered. To incorporate selection and MAF threshold, we consider the A locus as the causal variant under selection and B locus as the molecular marker under the MAF filtering. Four possible haplotypes (i.e., *A*_1_ *B*_1_, *A*_1_*B*_2_, *A*_2_*B*_*1*_, and *A*_2_*B*_2_) are possible at these two loci. In a population of size *N*_*e*_, the frequency counts of these four haplotypes can take on 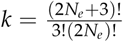 possible values, where for each of these possibilities, the sum of the four counts is 2*N*_*e*_. For example, when *N*_*e*_ is equal to two, all possible combinations of haplotype frequency counts are given in Figure 1.

**Figure 1.**
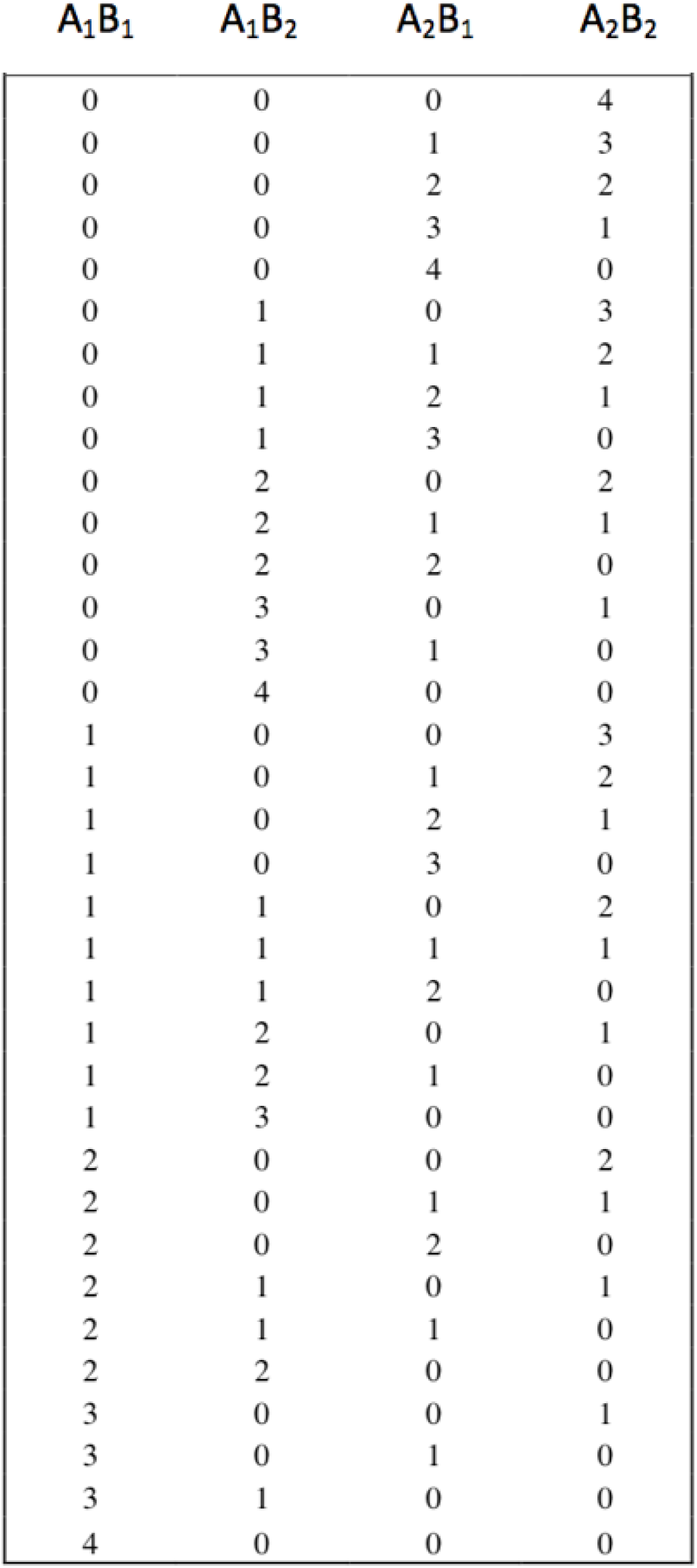
All possible combinations of haplotype frequency counts for a diploid population when *N*_*e*_ = 2

In general, let **X** be a *k* × 4 matrix with each row representing a possible combination of haplotype frequency counts for some value of *N*_*e*_. Thus, the rows of **X** represent the collection of k lines with different allele frequencies at two loci. Let **P**_*t*_ denote a *k* × 1 vector with element *i* indicating the probability of the frequency counts in row *i* of **X** at generation *t*. Thus, **P**_*t*_ gives the distribution of allele frequencies at generation *t*. We show below that the distribution in generation *t* + 1 can be written in general as

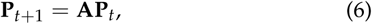

where **A** is a *k* × *k* transition matrix that is derived below in various circumstances of recombination, mutation and selection.

First, consider a line with haplotype frequency counts 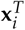 from *i*th row of **X** at generation *t*, e.g., 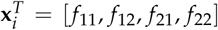, where the frequency counts for the four haplotypes, *A*_1_*B*_1_, *A*_1_*B*_2_, *A*_2_*B*_1_, and *A*_2_*B*_2_, are denoted by *f*_11_,*f*_12_,*f*_21_ and *f*_22_. Ignoring the effects of recombination, mutation, or selection, sampling of 2*N*_*e*_ gametes from this line can be modeled by a multinomial process with sample size *n* = 2*N*_*e*_ and probability vector 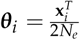 for the four haplotype probabilities. Thus, the distribution of the frequency count in the next generation is given by the *k* x 1 vector **m**_*i*_, where the element *j* of **m**_*i*_ is the probability of getting 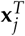 from the Multinomial(*n*, ***θ***_*i*_) distribution. Given that element *i* of **P**_*t*_ is the probability of 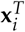 in generation *t*, the distribution of allele frequencies in the next generation is given by:

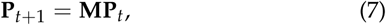

where **M** is the *k* × *k* matrix with columns m_*i*_, as described above, for *i* = 1,…,*k*. The above formula shows how the distribution of allele frequencies change due to genetic drift, ignoring recombination, mutation, and selection.

Now we accommodate recombination in computing **P**_*t*+1_. In gametes produced from generation *t*, the probability of a nonrecombinant *A*_1_*B*_1_ is 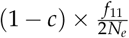, where *c* is the recombination rate between locus *A* and locus *B*. On the other hand, a recombinant *A*_1_ *B*_1_ can be produced in one of four ways. They and their associated probabilities are:

1. Alleles *A*_1_ and *B*_1_ originate from two different *A*_1_ *B*_1_ haplotypes with probability 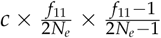;
2. Allele *A*_1_ originates from an *A*_1_ *B*_1_ haplotype and *B*_1_ originates from an *A*_1_ *B*_2_ haplotype with probability 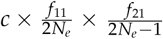;
3. Allele *A*_1_ originates from an *A*_1_ *B*_2_ haplotype and *B*_1_originates from an *A*_1_ *B*_1_ haplotype with probability 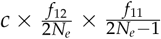;
4. Allele *A*_1_ originates from an *A*_1_ *B*_2_ haplotype and *B*_1_ originates from an *A*_2_ *B*_1_ haplotype with probability 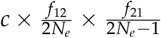.

Thus, accounting for recombination, the probability of a *A*_1_ *B*_1_ haplotype changes from 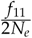 to:

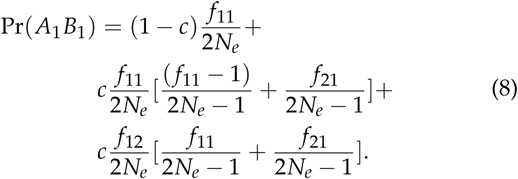

Similarly, due to recombination, the probabilities of the other three haplotypes become:

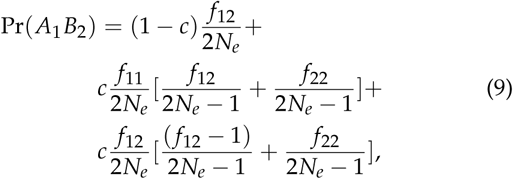

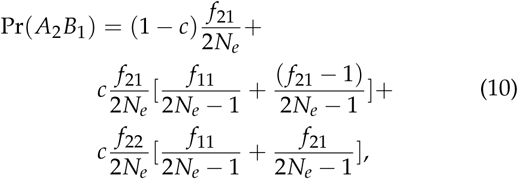

and

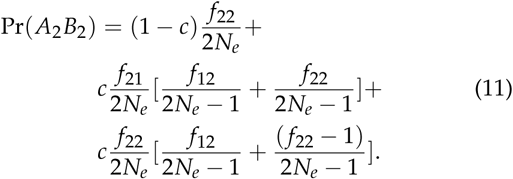

Now, to see how mutation alters these probabilities, we let 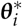 be a vector of the four probabilities from equations (8) through (11) computed using the four haplotype frequencies in 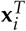. In modeling mutation, we assume that an *A*_1_ or *B*_1_ allele mutates to an *A*_2_ or *B*_2_ allele with probability *u* and an *A*_2_ or *B*_2_ allele mutates to an *A*_1_ or *B*_1_ allele with probability *v*. Then, haplotype probabilities following mutation can be computed as:

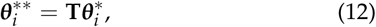

where

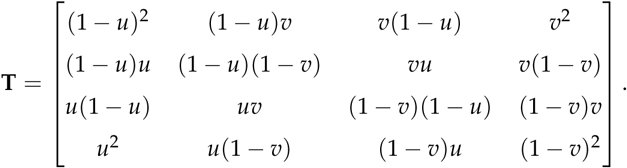

Furthermore, to incorporate selection in the model, a selection coefficient s that reduces the allele frequency of *A*_1_ at locus *A*, which is a causal variant, is considered. Conditional on the haplotype probabilities in 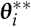, i.e., 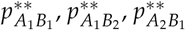 and 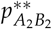, the haplotype probabilities following selection can be computed as

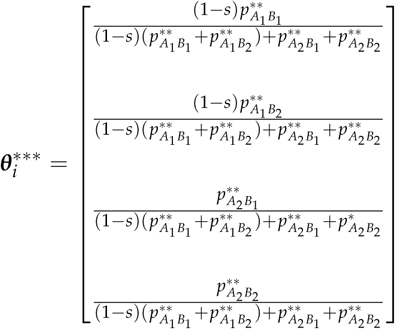

Finally, to compute the distribution of allele frequencies in generation *t* + 1 using Equation (6), where the forces of the genetic drift, recombination, mutation and selection are simultaneously considered, the matrix **A** is defined such that element *j* of column *i* contains the probability of getting 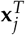 from a Multinomial 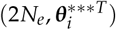 distribution, for *i, j* =1,…,*k*.

The value of *r*^2^ can be computed for a sub-population or line with haplotype frequency counts in any row of **X**. For example, consider the frequency counts in row 11 of the **X** matrix given in Figure 1, where *f*_11_ = 0, *f*_12_ = 2, *f*_21_= 1 and *f*_22_ = 1. From these frequency counts, *Pr*(*A*_1_) = 1/2, *Pr*(*B*_1_) = 1/4 and *Pr*(*A*_1_ *B*_1_) = 0 are obtained, and *r*^2^ calculated using equations (1) and (2) is 1/3 for a line with haplotype frequency counts in 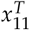 from Figure 1. The distribution of *r*^2^ in generation *t* is given by values of *r*^2^ corresponding to the frequency counts in each row of **X** together with the probabilities for haplotype frequency counts given in **P**_*t*_. Note that haplotype frequency counts in rows of **X** that have indeterminate values of *r*^2^ (i.e. when one or other allele is extinct) are not used to compute the distribution of *r*^2^. Similarly, the distribution of *r*^2^ with MAF cutoff is given by considering only the values of *r*^2^ and their probabilities corresponding to the rows of **X** with MAF at the B locus ≥ 5%. MAF of 0.05 is used as a threshold in the present study.

Starting with an allele frequency of 0.5 at each locus and linkage equilibrium between the two loci, the expected value of *r*^2^ was computed over generations given some values of the mutation rate, recombination rate, selection coefficient and effective population size. Mutation rate of *u* = *v* = 0 is used to represent the absence of mutation, while *u* = *v* = 1 × 10^−9^ is used to represent the existence of mutation. Similarly, a selection coefficient of *s* = 0 is used to represent the absence of selection, while s = 0.1 or s = 0.01 is used to represent the existence of selection.

### Data Analysis

In our analysis, different population parameters are considered. Populations with N_e_ ranging from 5 to 50 in intervals of 5 are compared under different recombination rates, mutation rates (i.e., *u* = 0 or *u* = 1*e*^−9^) and selection coefficients for the causal variant (i.e., *s* = 0.1 or *s* = 0.01 for locus *A*), in the absence or presence of filtering by MAF (MAF ≥ 0.05 for locus *B*). The recombination rates ranged from 0.01 to 0.1 in intervals of 0.01 and from 0.1 to 0.5 in intervals of 0.1.

The recombination rate for adjacent markers in a bovine 777k SNP panel was considered to mimic a realistic scenario. The recombination rate for adjacent markers was estimated to be 6.25 ×10^−5^ using the Kosambi map function given that the mean distance between adjacent markers is around 5 kb (Espigolan *et al*. 2013), and the average genetic distance per unit of physical distance in bovine genome is 1.25 cM/Mb (Arias *et al.* 2009).

A period of more than 3,000 generations was used to ensure that the haplotype frequencies had reached their equilibrium values. At equilibrium, the distribution of LD stays constant over generations (i.e., *P*_*t*+1_ = *P*_*t*_). That distribution was used to describe, quantify and compare 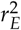 at different equilibria in the absence or presence of mutation, selection or MAF filtering. Results from a population similar to the international black and white Holstein dairy cattle are presented in these cases, for which the effective population size is estimated to be about 50 (Kim and Kirkpatrick 2009; Wray *et al*. 2019).

Furthermore, the exact distributions of 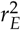 derived under different scenarios were used to validate the approximate deterministic formulas from Sved (Sved and Feldman 1973) and Hill (Hill 1975). To generalize our results to larger effective population size, a non-linear regression model of the form 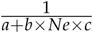, following Sved’s formula, was considered. The parameters in that model were estimated by non-linear least squares using the data generated from our transition-matrix approach.

The authors state that all data necessary for confirming the conclusions presented in the article are represented fully within the article.

## Results

### Comparison of Sved’s and Hill’s approximations to the exact distributions of 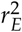 derived from the transition-matrix approach

In this section, Sved’s and Hill’s approximations were compared to the exact distribution of 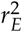 derived from our transition-matrix approach. The relationship between equilibrium value of 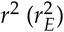, recombination rate and mutation rate is shown in Figure 2 for these three approaches. In the absence of mutation and selection, Sved’s formula showed consistency with the exact values from our transition-matrix approach (Figure 2(a)). On the other hand, Hill’s formula had a better fit to the exact values of 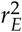 (Figure 2(b)) in the presence of mutation. However, neither Sved’s nor Hill’s deterministic formulae are accurate enough to describe 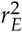 in the presence of mutation. Therefore, we proposed an adjusted non-linear regression model to correct Sved’s approximation to predict 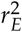 for larger effective population sizes in the presence of mutation. The mean square of errors is used to evaluate the fit of these three models and shows that our calibrated non-linear regression model was significantly better than Sved’s or Hill’s approximations.

**Figure 2.**
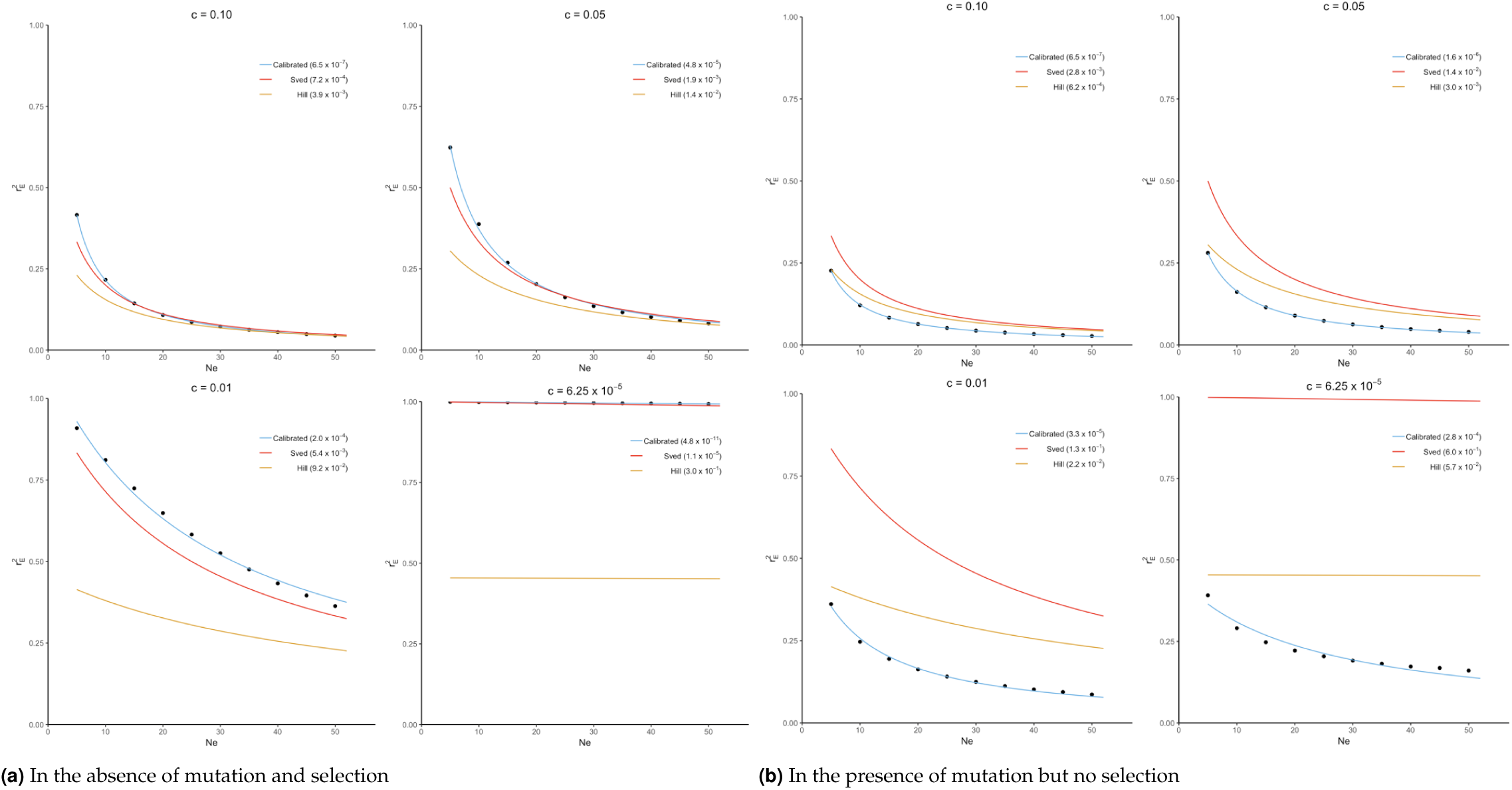
Comparison of Sved’s and Hill’s approximation to exact distribution of 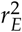 (scatter points) derived from transition-matrix approach. Mean square errors are shown in the parentheses. “Calibrated” denotes the non-linear regression formula derived from the transition-matrix approach

### Trajectory of LD over the generations until equilibrium is reached

The trajectory of evolving LD over generations computed from our transition-matrix approach is shown in this section, in the absence or presence of mutation and selection, where selection was applied with either a low selection coefficient of 0.01 or a high selection coefficient of 0.1. Starting with an allele frequency of 0.5 at each locus and linkage equilibrium between the two loci, expected values of LD (i.e., *E(r*^*2*^)) over generations are displayed in Figure 3. Note that the trajectory may be quite different, depending on the initial conditions used.

**Figure 3.**
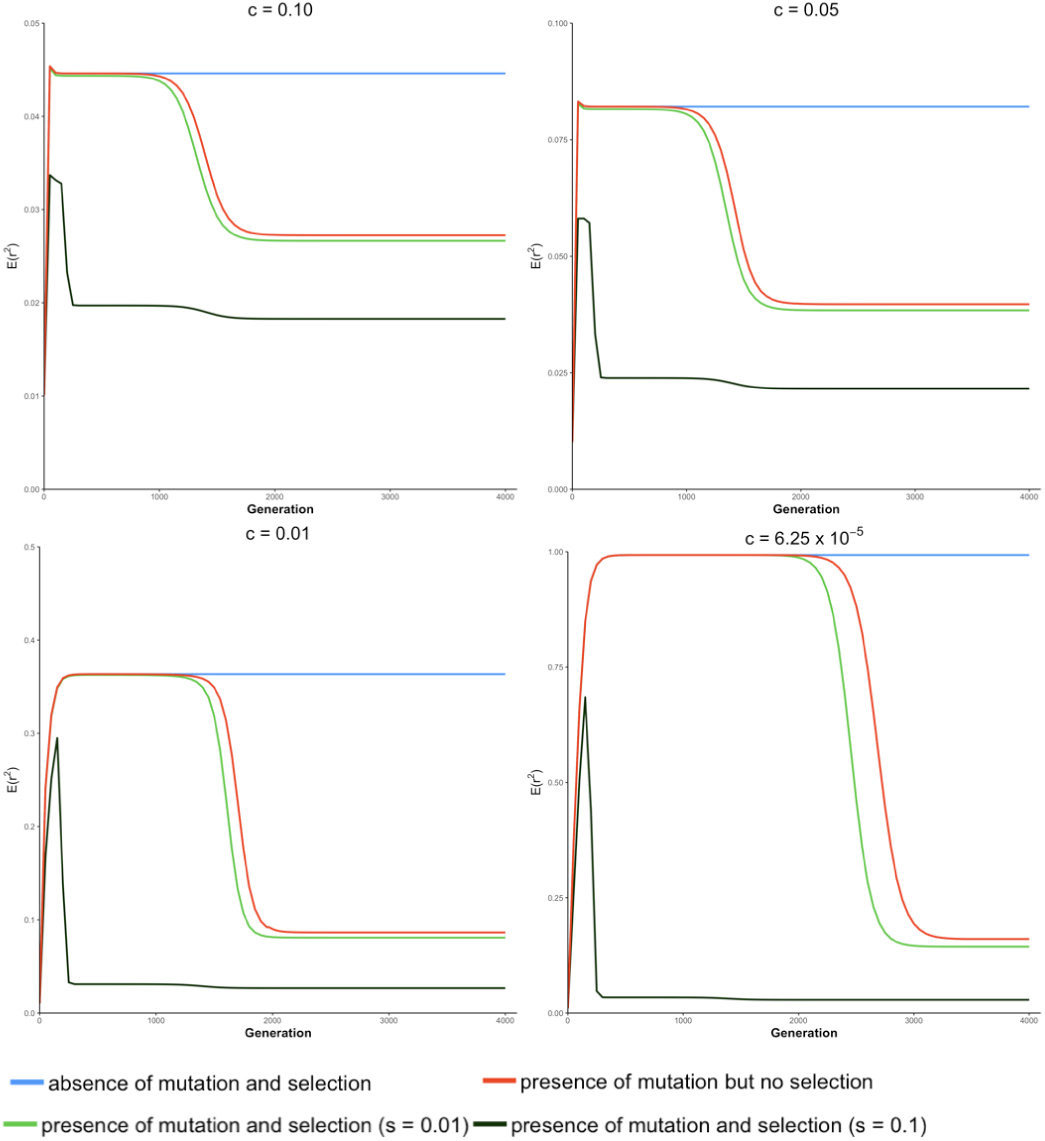
Expected value of LD (*E*(*r*^2^)) over generations under four different conditions at different recombination rates (c) for a population of effective size *Ne* = 50.

In the absence of mutation and selection, the expected value of *r*^2^ over generations increases to its maximum and reaches its “equilibrium” status. Value of 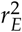 increases as c decreases due to the decreasing breakdown of LD with lower recombination rates. Typically, when the recombination rate is relatively small (e.g., *c* = 6.25 × 10^−5^), the expected value of LD at “equilibrium” is almost equal to 1. However, at this “equilibrium” stage, frequencies of lines (*P*_*t*_) keep changing though the distribution of allele frequencies remain unchanged. This is because forces of recombination and drift are balanced in lines with determinate *r*^2^, and frequencies of lines with determinate *r*^2^ almost proportionally decrease toward 0.

In the absence of selection but with mutation, the expected value of *r*^2^ increases and stays at an apparent equilibrium for several generations and then decreases to its true equilibrium value, where the mutation-drift equilibrium is reached. During the apparent equilibrium that is initially observed, the expected value of *r*^2^ is identical to the equilibrium value in the absence of mutation and selection, where the forces of drift and recombination are balanced. When this stage is first reached, the effect of mutation is negligible because the mutation rate is low relative to the frequency of lines with segregating loci. The apparent equilibrium ends when the frequency is high for lines where one or both loci are fixed (*r*^2^ is indeterminate) and the frequency of lines with segregating loci becomes close to the rate of mutation. Then, alleles introduced due to mutation into these lines have a noticeable effect on the distribution of allele frequencies and expected LD changes. At the end of the apparent equilibrium, the most frequent lines have only one haplotype. In the lines with two haplotypes, which have low frequencies, *r*^2^ is either indeterminate or has value 1.0. When mutation introduces a third haplotype in a line where *r*^2^ is 1.0, it drops in value. Further, when mutation introduces a third haplotype in a line where *r*^2^ is indeterminate, the value of *r*^2^ will be close to zero. Thus, as mutation becomes noticeable, the expected value of LD decreases. When the true equilibrium is reached, the loss of variation by fixation is balanced by its replenishment by mutation. In other words, the frequency of lines where loci are segregating will remain non-zero. Therefore, when mutation is present, the probabilities of allele frequencies (*P*_*t*_) stay constant over generations when the true equilibrium is reached.

In the presence of both mutation and selection, lines with segregating loci become low in frequency at an earlier stage due to selection, particularly when the selection coefficient is large (e.g., s = 0.1), and thus, mutation starts to have a noticeable effect on allele frequency at an earlier stage in the presence of selection relative to when selection is absent. When selection coefficient is relatively large (e.g., s = 0.1), the expected value of *r*^2^ reaches its maximum value early and then decreases sharply to two plateaus. When the recombination rate is low, e.g., *c* ≤ 0.01, the difference between these two plateaus is small.

In the absence of selection, allele frequencies at two loci tend to be similar. However, when locus A is under selection, allele frequencies at these two loci tend to be different such that LD is lower than when selection is absent. This effect, however, is negligible when the selection coefficient is small (e.g., s = 0.01). Thus, a selection coefficient of 0.1 is used to present results when mutation and selection are present in the next section.

The expected values of *r*^2^ over generations in the presence of filtering by MAF (i.e., MAF ≥ 0.05 for locus B) are shown in the Appendix (Figure 6), where a similar pattern as for E(*r*^2^) over generations are observed.

**Figure 4.**
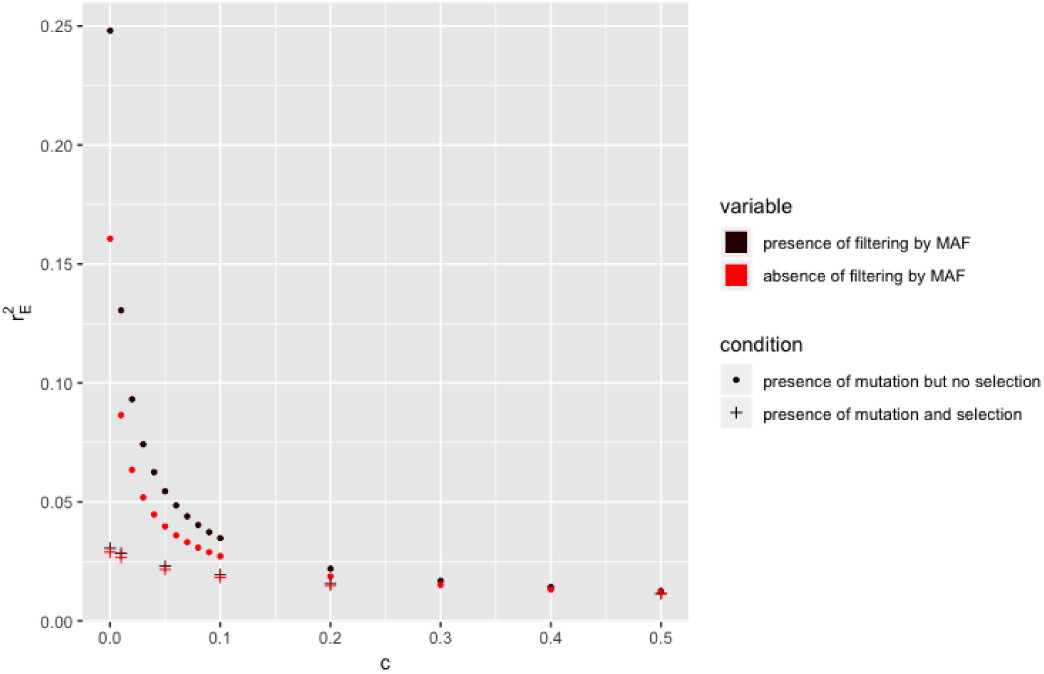
Relationship between the expectation of LD at equiLibrium 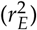 and recombination rate (*c*) in the absence or presence of selection or filtering by MAF at locus B for a population of effective size *N*_*e*_ = 50 with mutation rate *u =* 1.0 × 10^−9^. Only locus A is under selection with selection coefficient *s* = 0.1.

**Figure 5.**
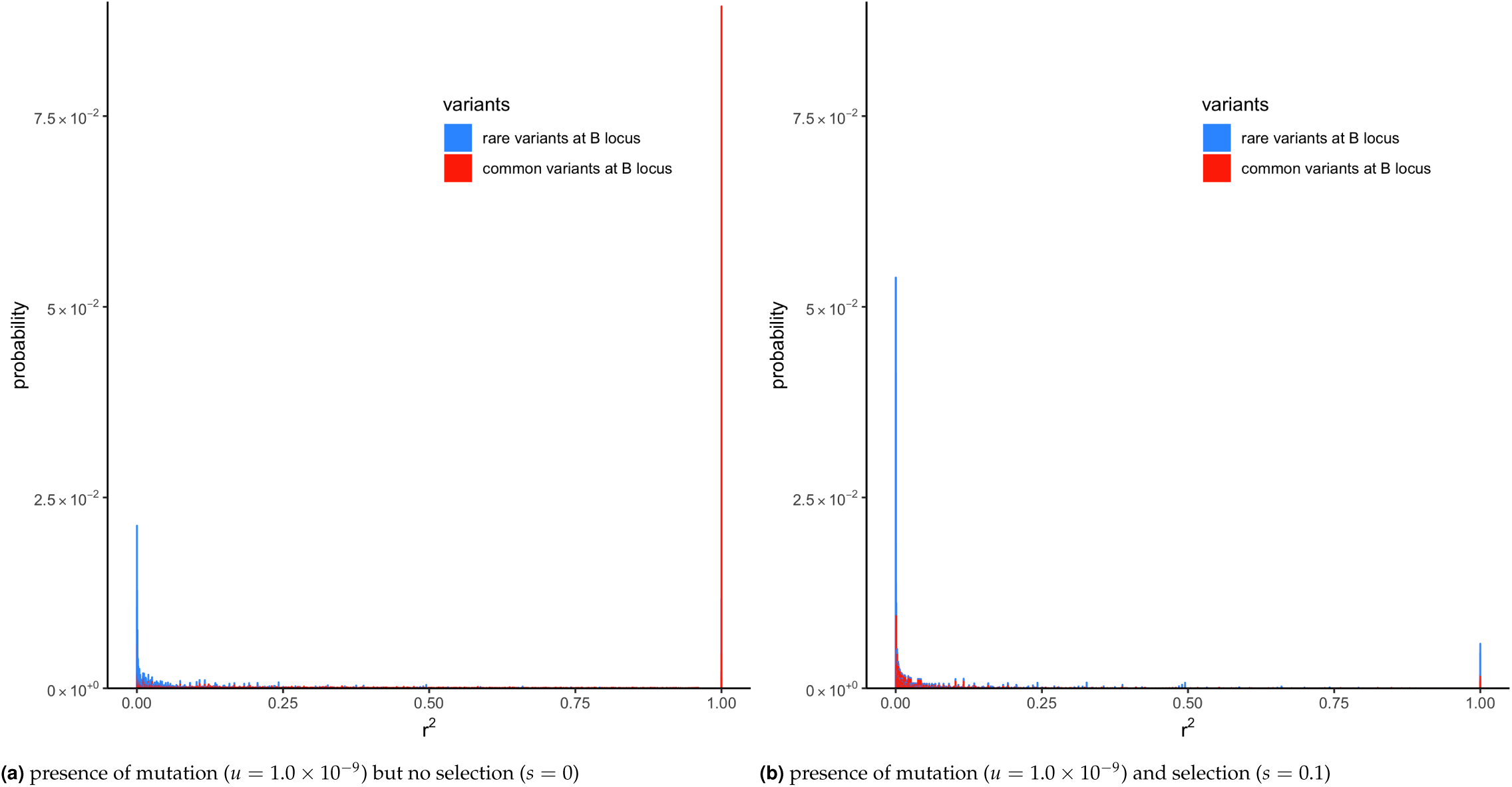
Distribution of LD (*r*^2^) with recombination rate (c) of 6.25 × 10 ^5^ at equilibrium in the presence of mutation but no selection or in the presence of mutation and selection in a population of effective size *Ne* = 50. Only locus A is under selection.

**Figure 6.**
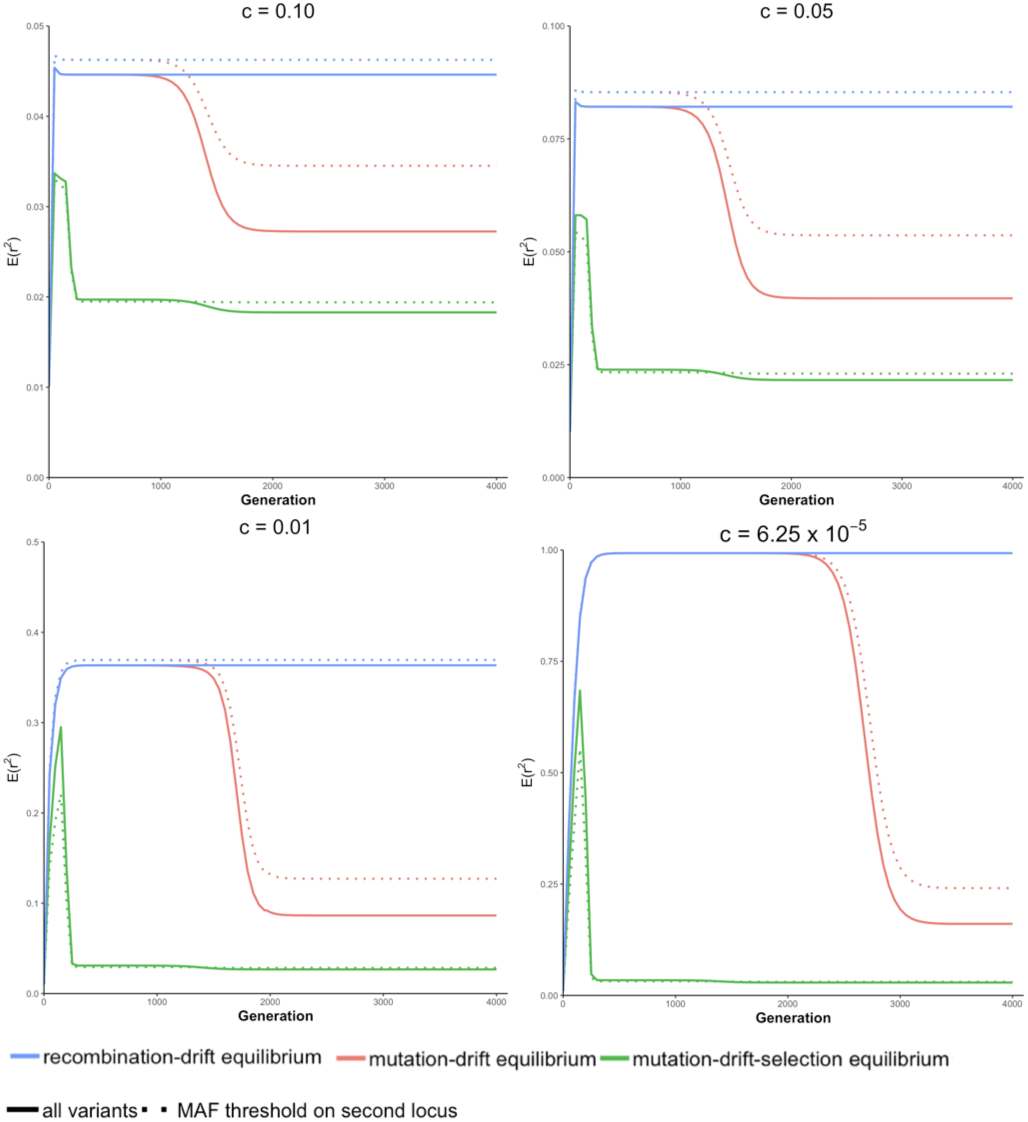
Expected value of LD (E(*r*^2^)) over generations in the absence or presence of filtering by MAF (i.e., MAF ≥ 0.05 for locus B) under three different conditions for a population of effective size *N*_*e*_ *=* 50.

### Distributions of LD at equilibrium

In a population where mutation is present and *N*_*e*_ = 50, the equlibrium value of 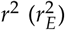 is shown in Figure 4, when selection is absent or present, for different recombination rates. As explained in last section, the equilibrium values of LD in the absence of selection are always higher than those in the presence of selection. The equilibrium value of LD (i.e., 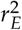) with filtering by MAF (i.e., MAF ≥ 5% at locus B) is higher than its corresponding value in the absence of filtering.

To understand why filtering by MAF increases 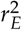, the *k* lines with different haplotype frequencies were divided into two groups: lines where MAF at the locus B is < 5% and lines where MAF ≥ 5%. Next, the transition-matrix approach was used to compute the frequency of each of these lines at equilibrium. In lines where *r*^2^ is defined, the equilibrium frequency of each line is plotted against its *r*^2^ value in Figure 5; the color blue is used for the first group of lines where MAF at the locus B is < 5% and red is used for the second group of lines where MAF is ≥ 5%; frequencies in the absence of selection are given in plot (a) and those in the presence of selection are given in plot (b).

In the absence of selection (plot (a)), a proportion of about 0.41 of the lines were in the first group and a proportion of about 0.59 were in the second group. It can be seen from this figure that most of the *r*^2^ values in the first group were small, where around 45% of the lines had an *r*^2^ value less than 0.001. In contrast, the second group had many lines with large *r*^2^ values; around 15% of the lines had *r*^2^ = 1. This difference in the distribution of *r*^2^ in the two groups shows why filtering by MAF at the B locus (removing lines from the first group, which had an abundance of low *r*^2^ values) would increase the expected value of *r*^2^.

The reason for the lower value of *r*^2^ in the first group is that, due to filtering by MAF at the B locus, this group has a large proportion of lines with recent mutations at the B locus. Consider a line where alleles are segregating at the A locus but fixed at the B locus. Such a line would belong to neither of the groups because *r*^2^ is not defined in a line where one locus is fixed. However, the introduction of a new allele at the B locus due to mutation will result in *r*^2^ becoming defined, but, typically, at a very low level because it results from the association in a single haplotype. The MAF in this line with the new mutation will be 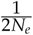, and it will belong to the first group provided that 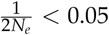.

Plot (b) of Figure 5 gives the distribution of *r*^2^ for the two groups in the presence of selection, and it can be seen that still there is a greater abundance of low *r*^2^ values in the first group. The difference between the two groups, however, is smaller than in the absence of selection, and this explains the smaller effect of filtering on expected *r*^2^ when selection is present (Figure 4).

### Extrapolation of exact 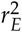 to larger population size by non-linear modelling

The two existing deterministic formulas (i.e., Sved’s and Hill’s formula) for 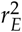 are not accurate as shown in Figure 2. Thus, a non-linear regression formula for recombination rate was calibrated using the exact values of 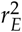 calculated from our transition-matrix approach. In the following, this formula is referred to as the calibrated non-linear regression formula. The form of the non-linear regression formula follows that of Sved’s formula as presented below:

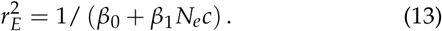

To study the extrapolation of 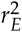 from small population sizes (i.e., *N*_*e*_ ≤ 50) to predict those for a larger population size, we split the values of 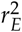 calculated from our transition-matrix approach into training and validation sets. Exact values of 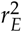 with *N*_*e*_ ≤ 40 are included in the training set, and the remaining with *N*_*e*_ > 40 are used for validation. The prediction accuracy is assessed using mean square error (MSE). As shown in Table 2 in the Appendix, prediction accuracy from the calibrated non-linear regression formula is substantially higher than those from Sved’s or Hill’s formulas. Finally, parameters in equation (13) are estimated using all available values of 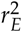, and parameter estimates are shown in Table 1. Typically, the estimates of *β*_0_ and *β*_1_ are inversely related to the values of recombination rates.

**Table 1.** Calibration (estimation of *β*_*i*_s) of the non-linear regression model under different recombination rates (c).

| c | $\beta_0$ | $\beta_1$ |
| --- | --- | --- |
| $6.25 \times 10^{-5}$ | 2.26 | 1557.11 |
| 0.01 | 1.75 | 21.41 |
| 0.02 | 1.49 | 15.13 |
| 0.03 | 1.31 | 12.58 |
| 0.04 | 1.17 | 11.10 |
| 0.05 | 1.05 | 10.09 |
| 0.06 | 0.96 | 9.33 |
| 0.07 | 0.87 | 8.74 |
| 0.08 | 0.80 | 8.25 |
| 0.09 | 0.73 | 7.84 |
| 0.1 | 0.67 | 7.49 |
| 0.2 | 0.28 | 5.48 |
| 0.3 | 0.02 | 4.49 |
| 0.4 | -0.19 | 3.85 |
| 0.5 | -0.38 | 3.38 |

**Table 2.** Comparisons of mean squared errors among the calibrated formula, Sved’s formula and Hill’s formula under different recombination rates (c) using validation data sets.

| c | Calibrated | Sved | Hill |
| --- | --- | --- | --- |
| $6.25 \times 10^{-5}$ | $7.39 \times 10^{-4}$ | $6.78 \times 10^{-1}$ | $8.31 \times 10^{-2}$ |
| 0.01 | $5.10 \times 10^{-5}$ | $7.01 \times 10^{-2}$ | $2.22 \times 10^{-2}$ |
| 0.02 | $1.31 \times 10^{-5}$ | $2.22 \times 10^{-2}$ | $9.46 \times 10^{-3}$ |
| 0.03 | $6.15 \times 10^{-6}$ | $1.01 \times 10^{-2}$ | $4.95 \times 10^{-3}$ |
| 0.04 | $3.86 \times 10^{-6}$ | $5.42 \times 10^{-3}$ | $2.91 \times 10^{-3}$ |
| 0.05 | $2.81 \times 10^{-6}$ | $3.24 \times 10^{-3}$ | $1.85 \times 10^{-3}$ |
| 0.06 | $2.21 \times 10^{-6}$ | $2.08 \times 10^{-3}$ | $1.24 \times 10^{-3}$ |
| 0.07 | $1.82 \times 10^{-6}$ | $1.41 \times 10^{-3}$ | $8.61 \times 10^{-4}$ |
| 0.08 | $1.53 \times 10^{-6}$ | $9.84 \times 10^{-4}$ | $6.17 \times 10^{-4}$ |
| 0.09 | $1.31 \times 10^{-6}$ | $7.09 \times 10^{-4}$ | $4.52 \times 10^{-4}$ |
| 0.1 | $1.13 \times 10^{-6}$ | $5.22 \times 10^{-4}$ | $3.37 \times 10^{-4}$ |
| 0.2 | $3.24 \times 10^{-7}$ | $4.11 \times 10^{-5}$ | $2.60 \times 10^{-5}$ |
| 0.3 | $1.13 \times 10^{-7}$ | $1.95 \times 10^{-6}$ | $6.04 \times 10^{-7}$ |
| 0.4 | $4.32 \times 10^{-8}$ | $9.39 \times 10^{-7}$ | $1.77 \times 10^{-6}$ |
| 0.5 | $1.68 \times 10^{-8}$ | $5.56 \times 10^{-6}$ | $6.73 \times 10^{-6}$ |

To further validate the accuracy of the extrapolation approach, the expected value of LD at equilibrium between adjacent markers in a 777k SNPs chip (i.e.,*c* = 6.25 × 10^—5^) was predicted for a Nellore cattle population, for which the effective population size is about 100 (Brito *et al*. 2013), using equation (13) with *β*_0_ = 2.26 and *β*_1_ = 1557.11. In a real genotyped Nellore cattle population (Espigolan *et al*. 2013), the estimated mean and standard deviation for 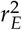 was 0.17 and 0.20, respectively. The predicted 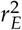 is 0.08, 0.449, and 0.976 using the calibrated non-linear regression formula, Hill’s formula, and Sved’s formula, respectively. Only the predicted 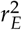 from the calibrated non-linear regression formula falls in the range 0.17 ± 0.2, while the other two are out of the measured range from this real population.

## Discussion

A contribution of this article is to propose a computational transition-matrix approach to deriving the distribution of LD between two loci over generations in the presence of multiple genetic forces including drift, mutation, and selection. These distributions of LD are also studied in the presence of filtering by MAF (i.e. MAF ≥ 0.05). In addition to deriving exact distribution of LD, several critical results emerge from the proposed approach.

1. **The expected value of LD at equilibrium decreases in the presence of selection.** Decrease of LD caused by mutation is magnified in the presence of selection due to a higher fixation rate of the favorable allele. In the presence of mutation but no selection (or in the presence of mutation and weak selection), allele frequencies at both loci are similar due to genetic drift, and the LD between them tends to remain high. Conversely, in the presence of mutation and strong selection, this phenomenon is disrupted by the selection process resulting in diverse gene frequencies at the two loci,and low LD bewteen them is observed.
2. **Caution is needed when LD between a causal variant and a marker is inferred after filtering out marker loci with low MAF.** LD between a causal variant and a marker variant with high MAF (i.e., MAF ≥ 5%) is higher than that between a causal variant and all marker variants, especially in the presence of mutation but no selection. That is, 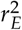 attributed to LD between a causal variant and marker variants with high MAF is relatively higher than that between a causal variant and marker variant with low MAF which leads to the reduction of the overall expected LD. In practice, LD is sometimes inferred from molecular marker panel with MAF cutoff applied. The inferred value is usually used to estimate important population parameters (e.g., effective population size). Here we have demonstrated the potential increase of LD brought by MAF cutoff and caution is needed when inferring LD in the presence of filtering by MAF.
3. **“Fake” equilibrium may appear in the presence of mutation.** Two equilibrium stages are observed in the presence of mutation. The first “equilibrium” indicates the balance between fixation increasing LD and recombination decreasing LD in lines with determinant *r*^2^ given the effect of mutation is negligible. When most loci become fixed at the end of the first “equilibrium” stage, change of allele frequency caused by mutation is of importance and reduces the expected LD. At the second equilibrium stage, the balance between mutation decreasing LD and fixation increasing LD is reached. Note that linkage equilibrium between two loci of allele frequency 0.5 is assumed in the initial population in Figure 3, and distribution of allele frequencies in the initial population affects the existence of the “fake” equilibrium.

### Exact 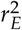 for large effective population sizes

The maximum effective population size (*N*_*e*_) presented in this paper is 50 (Kim and Kirkpatrick 2009; Wray *et al*. 2019), which is the estimated *N*_*e*_ for the international black and white Holstein dairy cattle population. Larger *N*_*e*_ was not studied in this paper due to computational limitations. When *N*_*e*_ = 50, the transition matrix **A** is square of order 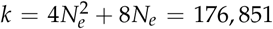, which requires more than 250 gigabytes memory to store. In addition, it takes around 5 hours to complete the computational process for the analysis of 3500 generations. Note that the memory complexity to store **A** is 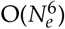. When *N*_*e*_ = 100, the estimated processing memory to store the transition matrix A is more than 14 terabytes. This computational problem may be addressed by parallel computing. This idea, however, needs further investigation and is out of the scope of this paper. Thus, to generalize the transition-matrix approach a non-linear regression model was calibrated using exact values of 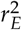. The proposed calibrations may provide a better description of the relationship between LD, effective population size, recombination rate,and mutation.

## Acknowledgments

This work was supported by the United States Department of Agriculture, Agriculture and Food Research Initiative National Institute of Food and Agriculture Competitive grant no. 2018-67015-27957. RLF is grateful to Prof. William G. Hill for extensive discussions of early results from the approach presented here.

